# Infant brain regional cerebral blood flow dynamics supporting emergence of vital functional networks

**DOI:** 10.1101/2021.02.08.430158

**Authors:** Qinlin Yu, Minhui Ouyang, John A. Detre, Huiying Kang, Di Hu, Bo Hong, Fang Fang, Yun Peng, Hao Huang

## Abstract

Human infancy is characterized by most rapid cerebral blood flow (rCBF) increases across lifespan and emergence of a fundamental brain system default-mode network (DMN). However, how infant rCBF change spatiotemporally across the brain and how the rCBF dynamics support emergence of vital functional networks such as DMN remains unknown. Here, by acquiring cutting-edge multi-modal MRI including pseudo-continuous arterial-spin-labeled perfusion MRI and resting-state functional MRI of infants aged 0 to 24 months, we elucidated unprecedented 4D spatiotemporal infant rCBF framework and region-specific physiology-function coupling across infancy. We found faster rCBF increases in the DMN than other regions. We also found strongly coupled increases of rCBF and network strength specifically in the DMN, suggesting faster local blood flow increase to meet extra neuronal metabolic demands in the DMN maturation. These results offer insights into physiological mechanism of brain functional network emergence and have important implications in altered network maturation in brain disorders.

## INTRODUCTION

The adult human brain receives 15-20% of cardiac output despite only representing 2% of body mass (Bouma and Muizelaar, 1990; Satterthwaite et al., 2014). Vast energy demand from the human brain starts from infancy which is characterized by fastest energy expenditure increase across lifespan (Pontzer et al., 2021). Infancy is also the most dynamic phase of brain development across entire lifespan with fastest functional and structural brain development. For example, during infancy the brain size increases dramatically in parallel with rapid elaboration of new synapses, reaching 80-90% of lifetime maximum by age of year 2 (Knickmeyer et al., 2008; Pfefferbaum et al., 1994; Ouyang et al., 2019a). Structural and functional changes of infant brain are underlaid by rapid and precisely regulated (Huang et al., 2013; Silbereis et al., 2016) spatiotemporal cellular and molecular processes including neurogenesis and neuronal migration (Rakic, 1995; Sidman and Rakic, 1973), synaptic formation (Huttenlocher and Dabholkar, 1997), dendritic arborization (Bystron et al., 2008; Ouyang et al., 2019b), axonal growth (Haynes et al., 2005; Innocenti and Price, 2005) and myelination (Miller et al., 2012; Yakovlev, 1967). These developmental processes demand rapidly increasing feed of energy to fuel the brain development. However, there have been no known whole-brain mappings of heterogeneous infant brain regional cerebral blood flow (rCBF) dynamics across landmark infant ages thus far, impeding fundamental view of energy expenditure across functional systems of early developing brain. As a result of differential neuronal growth across cortex, functional networks in the human brain develop differentially following the order from primary sensorimotor to higher-order cognitive systems (Cao et al., 2017a; Huang and Vasung, 2014; Tau and Peterson, 2010; Yu et al., 2016). The default-mode network (DMN) (Raichle et al., 2001) is widely recognized as a fundamental neurobiological system associated with cognitive processes that are directed toward the self and has important implication in typical and atypical brain development (Buckner et al., 2008). Unlike primary sensorimotor (SM) and visual (Vis) networks emerging relatively earlier around and before birth (Cao et al., 2017b; Doria et al., 2010; Smyser et al., 2010), emergence of the vital resting-state DMN is not well established until late infancy (Gao et al., 2009). Till now, it has been unclear how emergence of vital functional networks such as DMN is coupled with rCBF increase during infancy.

Regional brain metabolism, including glucose utilization and oxygen consumption, is closely coupled to regional CBF (rCBF) that delivers the glucose and oxygen needed to sustain metabolic needs (Raichle et al., 2001; Vaishnavi et al., 2010). Infant rCBF has been conventionally measured with positron emission tomography (PET) (Chugani and Phelps, 1986; Chugani et al., 1987) and single-photon emission computerized tomography (SPECT) (Chiron et al., 1992) which are not applicable to infants due to the associated exposure to radioactive tracers. By labeling the blood in internal carotid and vertebral arteries in neck and measuring downstream labeled arterial blood in brain, arterial-spin-labeled (ASL) (Alsop et al., 2015; Detre and Alsop, 1999) perfusion MRI provides a method for noninvasive quantifying rCBF without requiring radioactive tracers or exogenous contrast agents. Accordingly, ASL is especially suitable for rCBF measurements of infants (Ouyang et al., 2017; Wang et al., 2008) and children (Jain et al., 2012; Satterthwaite et al., 2014). Phase-contrast (PC) MRI, utilizing the phase shift proportional to velocity of the blood spins, has also been used to measure global CBF of the entire brain (Liu et al., 2019). Through integration of pseudo-continuous ASL (pCASL) and PC MRI, rCBF measured from pCASL can be calibrated by global CBF from PC MRI for more accurate infant brain rCBF measurement (Aslan et al., 2010; Ouyang et al., 2017). With rCBF closely related to regional cerebral metabolic rate of oxygen (CMRO2) and glucose (CMRGlu) at the resting state in human brain (Fox and Raichle, 1986; Gur et al., 2009; Paulson et al., 2010; Vaishnavi et al., 2010), rCBF could be used as a surrogate measure of local cerebral metabolic level for resting infant brains.

Early developing human brain functional networks can be reproducibly measured with resting-state fMRI (rs-fMRI). For example, a large scale of functional architecture at birth (Cao et al., 2017b; Doria et al., 2010; Fransson et al., 2008) has been revealed with rs-fMRI. Functional networks consist of densely linked hub regions to support efficient neuronal signaling and communication. These hub regions can be delineated with data-driven independent component analysis (ICA) of rs-fMRI data and serve as functional regions-of-interests (ROIs) for testing physiology-function relationship. Meeting metabolic demands in these hub ROIs is critical for functional network maturation. In fact, spatial correlation of rCBF to the functional connectivity (FC) in these functional ROIs was found in adult brains (Liang et al., 2013). Altered DMN plays a vital role in neurodevelopmental disorders such as autism (Doyle-Thomas et al., 2015; Lynch et al., 2013; Padmanabhan et al., 2017; Washington et al., 2014). Thus, understanding physiological underpinning of the DMN maturation offers invaluable insights into the mechanism of typical and atypical brain development.

We hypothesized heterogeneous rCBF maps at landmark infant ages and faster rCBF increase in brain regions of higher cognitive functions (namely DMN regions) during infancy than those of primary sensorimotor functions where functional networks emerge before or around birth (Cao et al., 2017a; Cao et al., 2017b; Doria et al., 2010; Fransson et al., 2007; Smyser et al., 2010; Peng et al., 2020). Furthermore, with rCBF an indicator of local metabolic level of glucose and oxygen consumption, we hypothesized strongly coupled rCBF and FC increases specifically in the DMN regions during infancy to meet extra metabolic demand of DMN maturation. In this study, we acquired multi-modal MRI including both pCASL perfusion MRI and rs-fMRI of 48 infants aged 0 to 24 months to quantify rCBF and FC, respectively. RCBF at the voxel level and in functional network ROIs were measured to test the hypothesis of spatiotemporally differential rCBF increases during infancy. Maturation of FC in the DMN was delineated. Correlation of FC increase and rCBF increase in the DMN ROIs was tested and further confirmed with data-driven permutation analysis, the latter of which was to examine if the coupling of rCBF and FC takes place only in the DMN during infancy.

## RESULTS

### Faster rCBF increases in the DMN hub regions during infant brain development

The labeling plane and imaging slices of pCASL perfusion MRI of a representative infant brain, reconstructed internal carotid and vertebral arteries, and four PC MR images of the target arteries are shown in Figure 1a. The rCBF maps of infant brains were calculated based on pCASL perfusion MRI and calibrated by PC MRI. As an overview, axial rCBF maps of typically developing brains at milestone ages of 1, 6, 12, 18 and 24 months are demonstrated in Figure 1b. High quality of the rCBF maps can be appreciated by a clear contrast between white matter and gray matter. A general increase of blood flow across the brain gray matter from birth to 2 years of age is readily observed. Heterogeneous rCBF distribution at a given infant age can be appreciated from these maps. For example, higher rCBF values in primary visual cortex compared to other brain regions are clear in younger infant at around 1 month. Figure 1b also demonstrates differential rCBF increases across brain regions. RCBF increases are prominent in the posterior cingulate cortex (PCC, a vital hub of DMN network), indicated by green arrows. On the other hand, rCBF in the visual cortex is already higher (indicated by blue arrows) than other brain regions in early infancy and increases slowly across infant development. The adopted pCASL protocol is highly reproducible with intraclass correlation coefficient (ICC) 0.8854 calculated from pCASL scans of a randomly selected infant subject aged 17.6 month, shown in Figure 1- figure supplement 1. As shown in Figure 1- figure supplement 2, three functional network ROIs including DMN, visual (Vis)network, and sensorimotor (SM) network were generated from infant rs-fMRI data. With rCBF measured at these functional network ROIs, Figure 1- figure supplement 3 quantitatively exhibits spatial inhomogeneity of rCBF distribution regardless of age. These quantitative measurements are consistent to the observation of heterogeneous rCBF distribution in Figure 1b. Specifically, as shown in Figure 1 – figure supplement 3, significant heterogeneity of rCBF was found across regions (*F*(6, 282) = 122.6, *p* < 10^−10^) with ANOVA test. With further paired *t*-test between regions, the highest and lowest rCBF was found in the Vis (82.1 ml/100g/min) and SM (49.1 ml/100g/min) regions, respectively (all *ts*(47) > 4.17, *p* < 0.05), while rCBF in the DMN (67.8 ml/100g/min) regions was in the middle (all *ts* (47) > 2.87, *p* < 0.05). Within the DMN, rCBF in the PCC (75.4 ml/100g/min) and LTC (72.0 ml/100g/min) regions were significantly higher than rCBF in the MPFC (60.7 ml/100g/min) (both *ts*(47) > 8.22, *p* < 0.05) and IPL regions (59.4 ml/100g/min) (both *ts*(47) > 7.87, *p* < 0.05).

**Figure 1.**
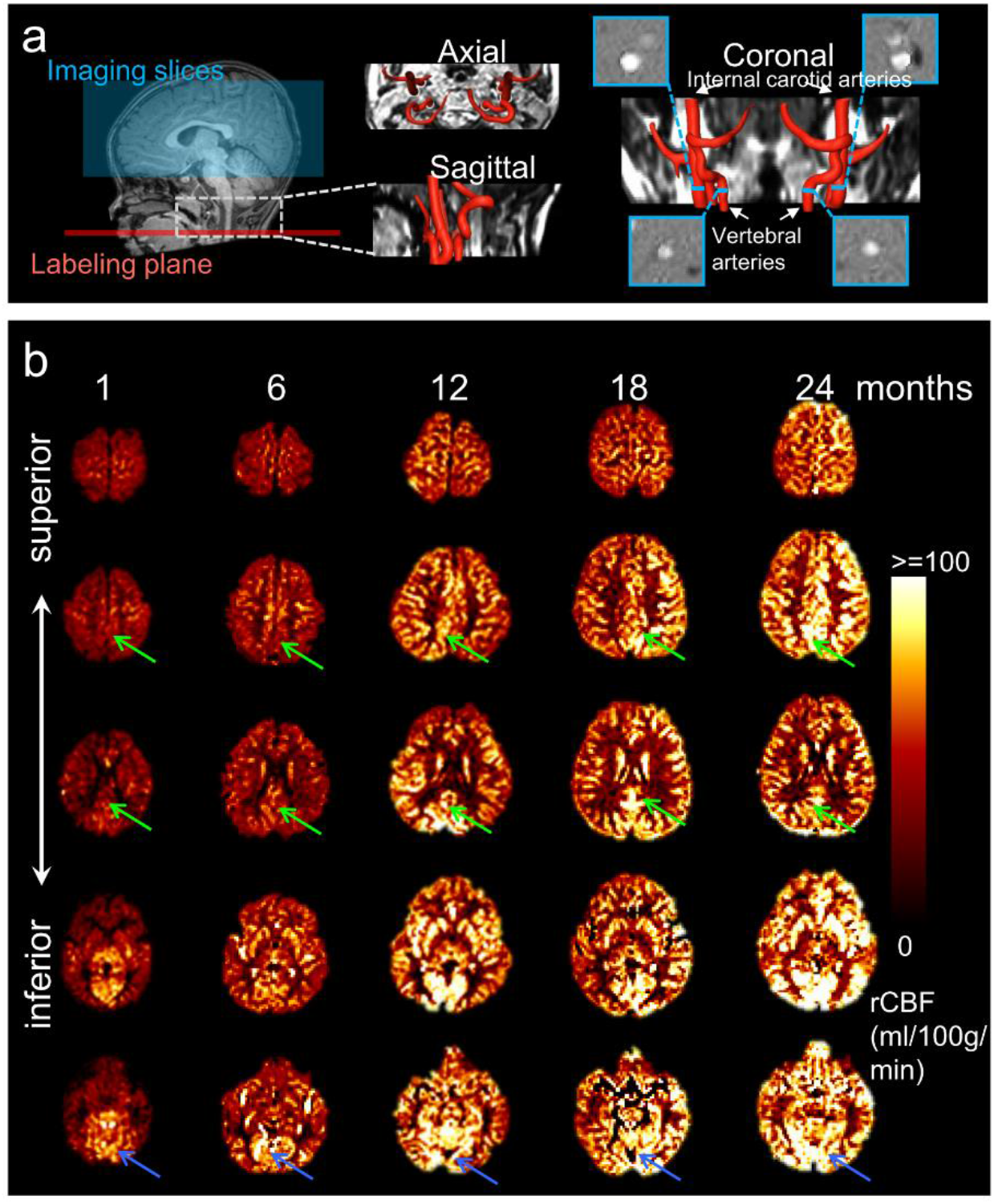
Acquisition of high-quality infant pseudo-continuous arterial-spin-labeled (pCASL) perfusion and phase contrast (PC) MRI and resultant axial regional cerebral blood flow (rCBF) maps at different infant ages. **a** Labeling plane (red line) and imaging volume (blue box) of pseudo-continuous arterial-spin-labeled perfusion MR imaging are shown on the mid-sagittal slice of T1-weighted image of a representative infant on the left panels. Axial and sagittal view of MR angiography with reconstructed internal carotid and vertebral arteries are shown in the middle of the panel a. On the right of the panel a, the coronal view of the reconstructed arteries is placed in the middle with four slices (shown as blue bars) of the PC MR scans positioned perpendicular to the respective feeding arteries. The PC MR images are shown on the four panels surrounding the coronal view of the angiography. These PC MR images measure the global cerebral blood flow of internal carotid and vertebral arteries and are used to calibrate rCBF. **b** rCBF maps of representative typically developing (TD) infant brains at 1, 6, 12, 18 and 24 months from left to right. Axial slices of rCBF maps from inferior to superior are shown from bottom to top of the panel b for each TD infant brain. Green arrows point to the posterior cingulate cortex (a hub of the DMN network) characterized by relatively lower rCBF at early infancy and prominent rCBF increases from 1 to 24 months. Blue arrows point to the visual cortex characterized by relatively higher rCBF at early infancy and relatively mild rCBF increase from 1 to 24 months. **Figure 1-figure supplement 1**. Highly reproducible pCASL protocol adopted in the present study for measuring rCBF. **Figure 1-figure supplement 2**. Identification of functional network regions-of-interests with resting-state fMRI. **Figure 1-figure supplement 3.** Heterogeneity of rCBF measurements across functional network regions.

Figure 2a shows cortical maps of linearly fitted rCBF values of infant brains from 0 to 24 months. Consistent with nonuniform profile of the rCBF maps observed in Figure 1b, the three-dimensionally reconstructed rCBF distribution maps in Figure 2a are also not uniform at each milestone infant age. RCBF increases from 0 to 24 months across cortical regions are apparent, as demonstrated by relatively high rCBF-age correlation r values across the cortical surface in Figure 2b. Heterogeneity of rCBF increases across all brain voxels can be more clearly appreciated in Figures 2a and 2b compared to Figure 1b. With DMN functional network regions including PCC, medial prefrontal cortex (MPFC), inferior posterior lobule (IPL) and lateral temporal cortex (LTC) as well as Vis and SM network regions delineated in Figure 1-figure supplement 2b as ROIs, rCBF trajectories in Figure 2c demonstrate that rCBF in these ROIs all increase significantly with age (Vis: *r* = 0.53, *p* < 10^−4^; SM: *r* = 0.52, *p* < 10^−4^; DMN: *r* = 0.7, *p* < 10^−7^; DMN_PCC: *r*= 0.66, *p* < 10^−6^; DMN_MPFC: *r* = 0.67, *p* < 10^−6^; DMN_IPL: *r* = 0.66, *p* < 10^−6^; DMN_LTC: *r* = 0.72, *p* < 10^−8^). Using the trajectory of primary sensorimotor (SM) (black line and circles) in Figure 2c as a reference, rCBF increase rates across functional network ROIs are also heterogeneous (Figure 2c). Specifically, significantly higher (all *p* < 0.05, FDR corrected) rCBF increase rate was found in total DMN ROIs (1.59 ml/100g/min/month) and individual DMN ROIs, including DMN_PCC (1.57 ml/100g/min/month), DMN_MPFC (1.63 ml/100g/min/month), DMN_IPL (1.42 ml/100g/min/month) and DMN_LTC (2.22 ml/100g/min/month), compared to in the SM ROI (0.85 ml/100g/min/month). Although the rCBF growth rate in the Vis (1.27 ml/100g/min/month) ROIs is higher than that in the SM ROIs, this difference is not significant (*p* = 0.13). Collectively, Figures 1 and 2 show that the CBF increases significantly and differentially across brain regions during infancy, with rCBF in the DMN hub regions increasing faster than rCBF in the SM and Vis regions (Figure 2). The 4D spatiotemporal whole-brain rCBF dynamics during infant development are presented in Video 1.

**Figure 2.**
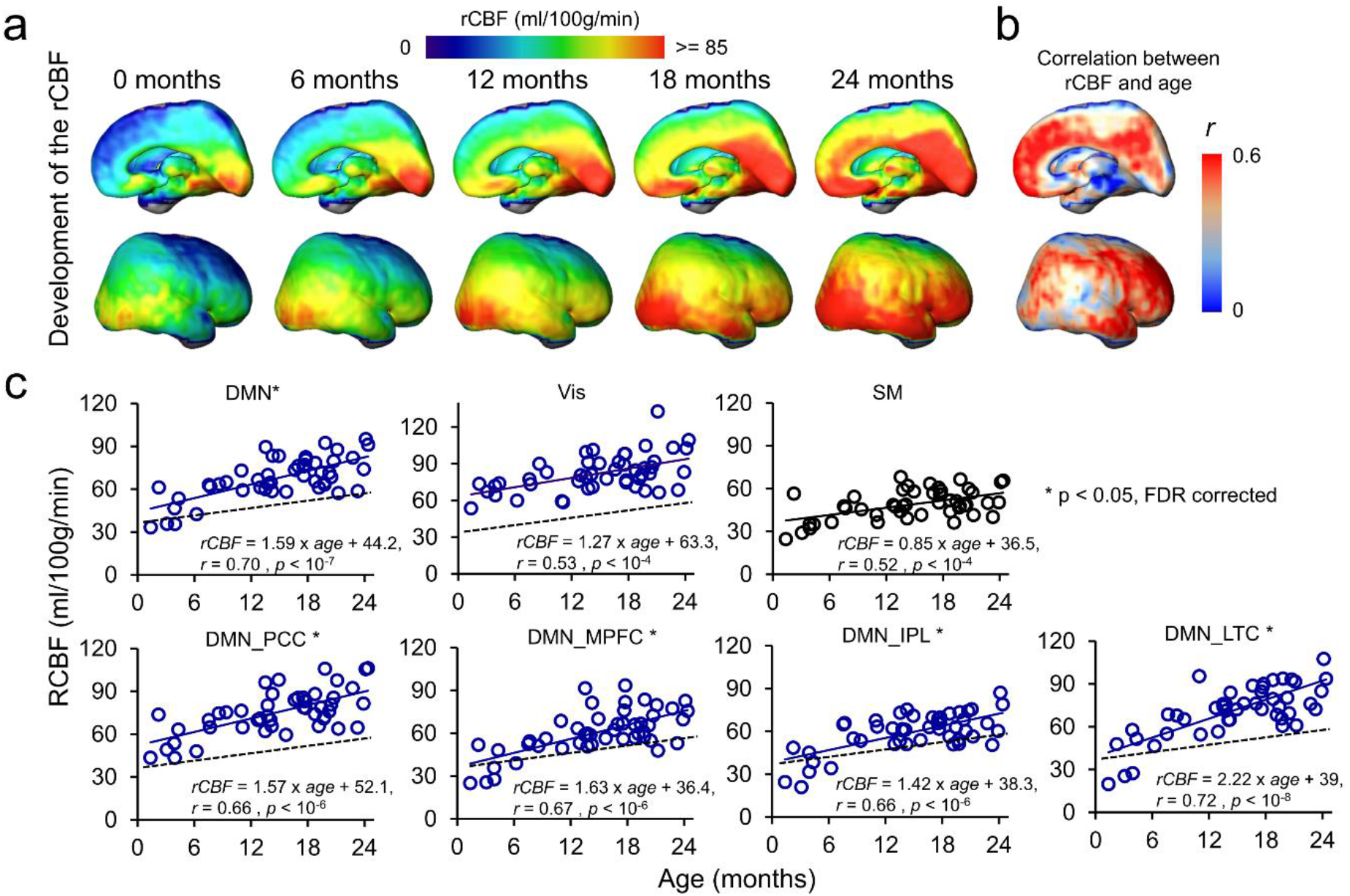
4D spatiotemporal regional cerebral blood flow (rCBF) dynamics and faster rCBF increases in the default-mode network (DMN) hub regions during infancy. **a** Medial (top row) and lateral (bottom row) views of fitted rCBF profiles of the infant brain at 0, 6, 12, 18 and 24 months in the custom-made infant template space demonstrate heterogeneous rCBF increase across the brain regions. **b** Medial (top) and lateral (bottom) views of rCBF-age correlation coefficient (r) map are demonstrated. **c** The scatterplots of rCBF measurements in the primary sensorimotor (SM) network (black circle and black line), visual (Vis) network (blue circle and blue line), and total and individual DMN hub regions (DMN_MPFC, DMN_PCC, DMN_IPL and DMN_LTC) (blue circle and blue line) demonstrate differential rCBF increase rates. * next to network name in each plot indicates significant (FDR-corrected *p*<0.05) differences of rCBF trajectory slopes from that of SM used as a reference and shown in a black dashed line. Abbreviations of DMN subregions: IPL: inferior posterior lobule; LTC: lateral temporal cortex; MPFC: medial prefrontal cortex; PCC: posterior cingulate cortex.

### Emergence of DMN during early brain development

Figure 3a shows emergence of the DMN in typically developing brain from 0 to 24 months as measured using rs-fMRI with a PCC seed region indicated by the black dash line. At around birth (0 months), the DMN is still immature with weak FC between other DMN regions (including MPFC, IPL, and LTC) and PCC (Figure 3a). During infant brain development from 0 to 24 months, Figure 3a shows that the functional connectivity between MPFC, IPL or ITC and PCC gradually strengthens. Figure 3b shows the FC-age correlation *r* value map. It can be appreciated from Figure 3b that across the cortical surface relatively higher r values are only located at the DMN regions (except the seed PCC).

**Figure 3.**
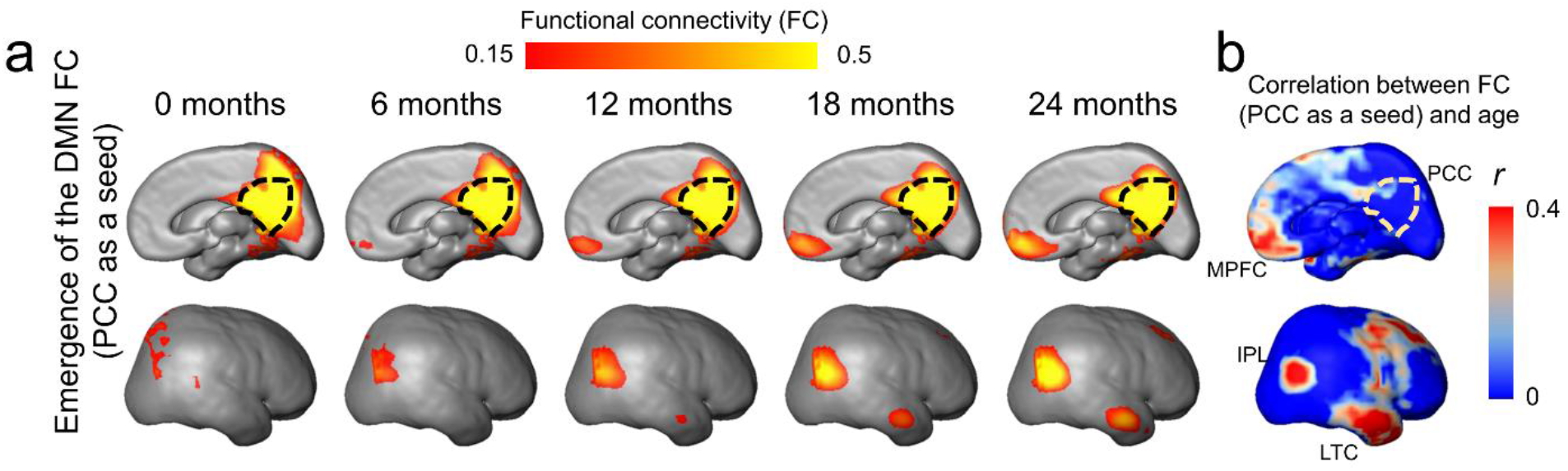
Emergence of functional connectivity (FC) within the default-mode network (DMN) during infancy. The maps of the DMN FC (posterior cingulate cortex (PCC) as a seed) at representative ages from 0 to 24 months are demonstrated in **a** and the map of correlation coefficient of FC (PCC as a seed) and age is demonstrated in **b**. In **a**, gradually emerging FC of other DMN regions (including MPFC, IPL and LTC) to the PCC from 0 to 24 months can be appreciated. The PCC is delineated by the black dashed contour. In **b**, stronger correlation between FC (PCC as a seed) and age is localized in DMN subregions IPL, ITL and MPFC. See legend of Figure 2 for abbreviations of the DMN subregions. **Figure 3-figure supplement 1**. Age-dependent changes of functional connectivity (FC) within the default-mode network (DMN), visual (Vis) and sensorimotor (SM) network regions during infancy.

As shown in Figure 3- figure supplement 1, only the FC within the DMN (*r* = 0.31, *p* < 0.05), but not the Vis (*r* = 0.048, *p* = 0.745) or SM (*r* = 0.087, *p* = 0.559), was found to increase significantly with age, indicating significant functional development in the DMN, but not in the Vis or SM network.

### Coupling between rCBF and FC within DMN during infant brain development

To test the hypothesis that rCBF increases in the DMN regions underlie emergence of this vital functional network, correlation between rCBF and FC was conducted across randomly selected voxels within the DMN of all infants aged 0-12 months (Figure 4a) and all infants aged 12-24 months (Figure 4b). Significant correlations (*p*<0.001) were found in both age groups. We further tested whether functional emergence of the DMN represented by increases of FC within the DMN (namely DMN FC) was correlated to rCBF increases specifically in the DMN regions, but not in primary sensorimotor (Vis or SM) regions. Figure 5a shows correlations between the DMN FC and rCBF at the DMN (red lines), Vis (green lines) or SM (blue lines) voxels. The correlations between the DMN FC and averaged rCBF in the DMN, Vis or SM region are represented by thickened lines in Figure 5a. A correlation map (Figure 5b) between the DMN FC and rCBF across the entire brain voxels was generated. The procedures of generating this correlation map are illustrated in Figure 5-figure supplement 1. The DMN, Vis and SM ROIs in Figure 5b were delineated with dashed red, green and blue contours, respectively, and obtained from Figure 1- figure supplement 2b. Most of significant correlations (r > r_crit_) between the DMN FC and voxel-wise rCBF were found in the voxels in the DMN regions, such as PCC, IPL and LTC, but not in the Vis or SM regions (Figure 5b). Demonstrated in a radar plot in Figure 5c, much higher percent of voxel with significant correlations between rCBF and the DMN FC was found in the DMN (36.7%, *p* < 0.0001) regions, than in the SM (14.6%, *p* > 0.05) or Vis (5.5%, *p* > 0.05) regions. Statistical significance of higher percent of voxels with significant correlations in the DMN (*p*<0.0001) was confirmed using nonparametric permutation tests with 10,000 permutations. We also conducted the correlation between the Vis FC and rCBF across the brain as well as permutation test. As expected, no significant correlation between the Vis FC and rCBF can be found in any voxel in the DMN, Vis or SM ROIs, demonstrated in Figure 5-figure supplement 2a. Similar analysis was also conducted for correlation between the SM FC and rCBF across the brain and percent of voxels with significant correlation was close to zero, as demonstrated in Figure 5-figure supplement 2b. Combined with the results shown in Figure 5, the results of coupling between Vis (Figure 5-figure supplement 2a) or SM (Figure 5-figure supplement 2b) FC and rCBF further demonstrated selected rCBF-FC coupling can be only found in the DMN ROIs, but not in the Vis or SM network ROIs.

**Figure 4.**
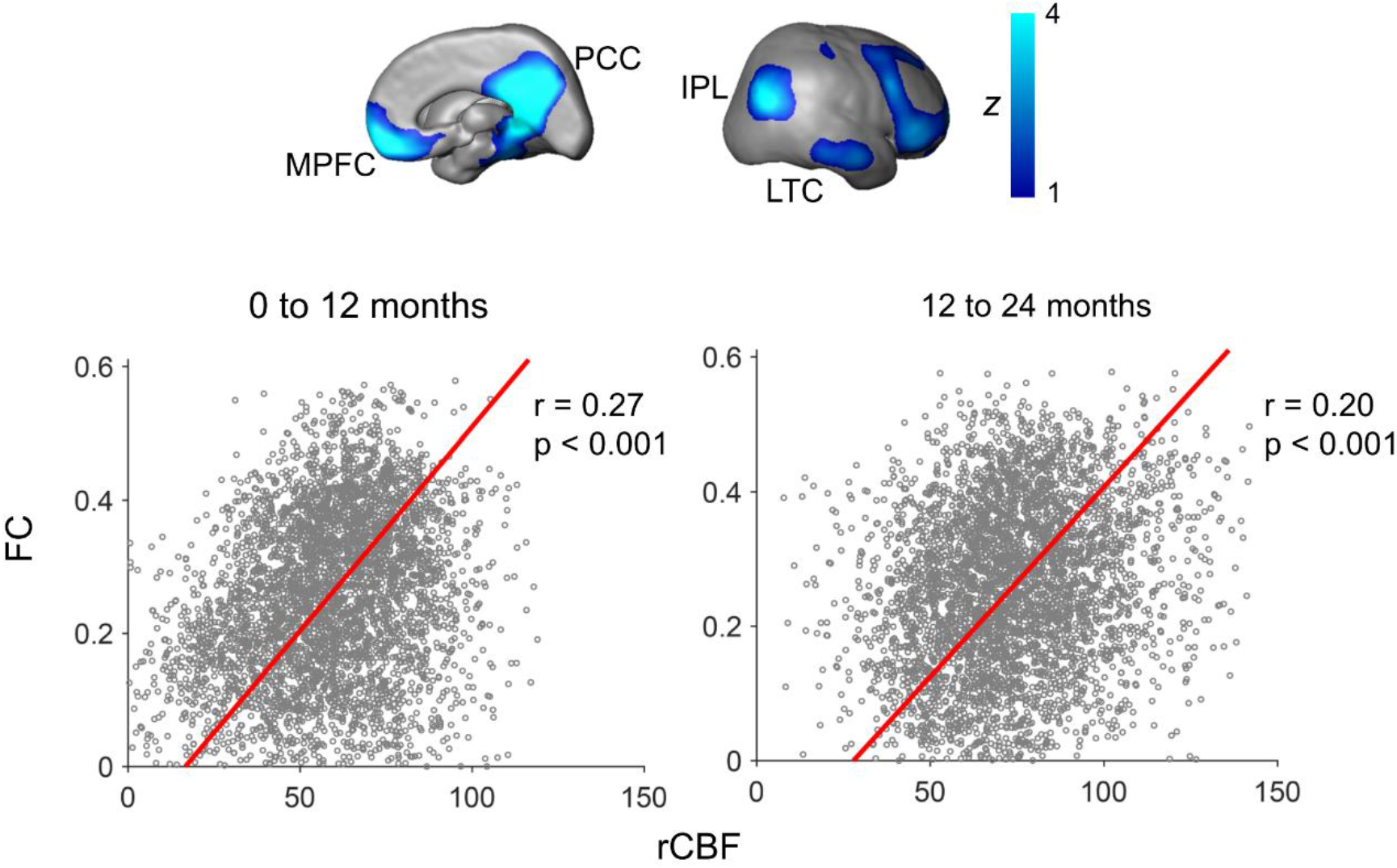
Significant correlation of regional cerebral blood flow (rCBF) and functional connectivity (FC) at randomly selected 4000 voxels within the default-mode network (DMN) for both infants aged 0-12 months (*p*<0.001, left scatter plot) and infants aged 12-24 months (p<0.001, right scatter plot). FC is the average of FC of a certain DMN voxel to all other DMN voxels. The DMN regions of interests obtained from a data-driven independent component analysis of resting-state fMRI of the 12-24-month infant cohort are shown on the top panels as an anatomical reference. See legend of Figure 2 for abbreviations of the DMN subregions.

**Figure 5.**
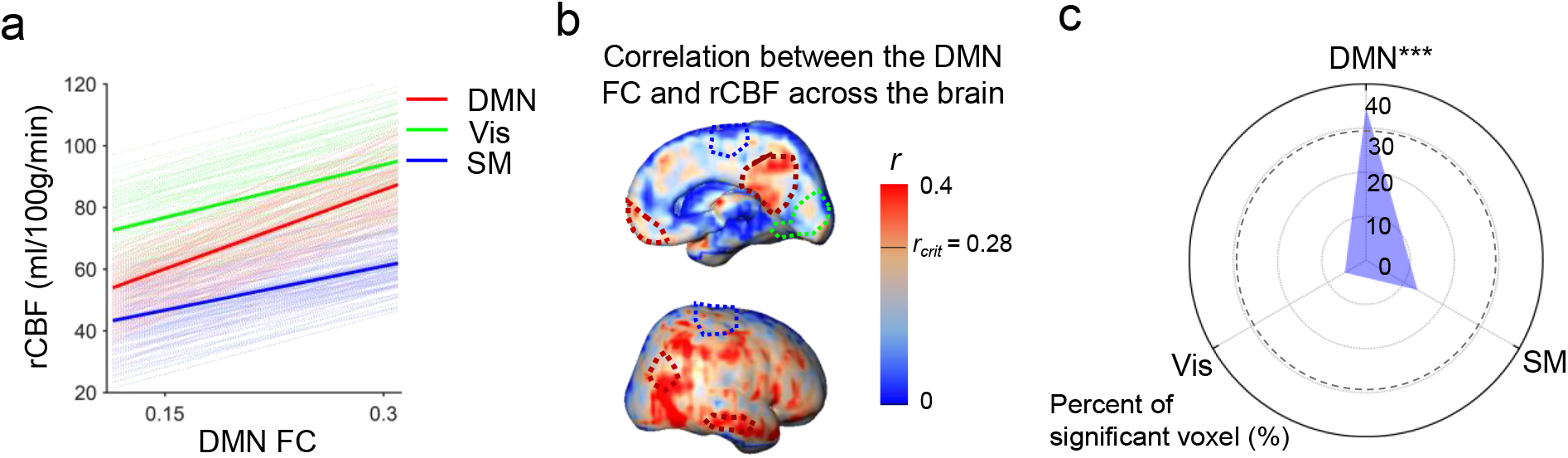
Significant correlation between functional emergence of the default-mode network (DMN) and regional cerebral blood flow (rCBF) increases specifically in the DMN regions, but not in primary sensorimotor (visual or sensorimotor) regions. **a** Correlation of intra-default-mode-network functional connectivity (DMN FC) and regional cerebral blood flow (rCBF) at randomly selected voxels in the DMN (light red lines), visual (Vis, light green lines) and sensorimotor (SM, light blue lines) network regions. Correlations of DMN FC and averaged rCBF in the DMN, Vis and SM network regions are shown as thickened red, green and blue lines, respectively. **b** Coupling between the DMN FC and rCBF across the brain can be appreciated by distribution of voxel-wise correlation coefficient (r) obtained from correlation between DMN FC and rCBF at each voxel. The short black line in the color bar indicates critical r value *r*_crit_ corresponding to *p*=0.05. Higher r values can be appreciated in the DMN hub regions including posterior cingulate cortex (PCC), medial prefrontal cortex (MPFC), inferior posterior lobule (IPL) and lateral temporal cortex (LTC) with their boundaries delineated by the dashed dark red contours (from Figure 1-figure supplement 2b). Dashed green and blue contours (also from Figure 1-figure supplement 2b) delineate the Vis and SM network regions, respectively. **c** Radar plot shows significant correlation between rCBF and intra-DMN FC in the DMN network (36.7%, *p* < 0.0001), but not in the Vis (14.6%, *p* > 0.05) or SM (5.5%, *p* > 0.05) networks. The radius represents the percent of the voxels with significant correlations between intra-DMN FC and rCBF in DMN, Vis and SM network regions, respectively. The dashed line circle indicates critical percent of significant voxels with *p*=0.05 from 10,000 permutation tests. *** indicates *p*<0.0001. **Figure 5-figure supplement 1**. Procedures of generating map of correlation between the default-mode network (DMN) functional connectivity (FC) and reginal cerebral blood flow (rCBF) across the entire brain voxels. **Figure 5-figure supplement 2**. Coupling between intra-visual network functional connectivity (Vis FC) and regional cerebral blood flow (rCBF) and coupling between intra-sensorimotor network (SM FC) and rCBF.

## DISCUSSION

We revealed strongly coupled rCBF and FC increases specifically in the DMN while establishing unprecedented 4D rCBF spatiotemporal dynamics during infancy. The tight rCBF-FC relationship found with multi-modal infant MRI suggests DMN emergence is supported by faster local blood flow increase in the DMN to meet metabolic demand, offering refreshing insight into the physiological mechanism underlying early brain functional architecture emergence. The delineated 4D brain perfusion spatiotemporal framework was characterized with heterogeneous rCBF distribution across brain regions at a specific age and differential age-dependent rCBF increase rates across brain regions during infant development, and can be used as quantified standard reference for detecting rCBF alterations (e.g. the z scores) of atypically developing brains. Elucidating the ontogeny of infant brain physiology and its functional correlates could greatly advance current understanding of general principles of early brain development.

Gradient of functional network maturations in early brain development has been more extensively characterized with recent rs-fMRI studies. Differential emergence of these functional networks is distinguished by different onset time as well as different maturational rate of various brain functions in a given developmental period. For example, primary sensory and motor functional networks, such as the SM and Vis networks, appear earlier before or around birth (Cao et al., 2017a; Doria et al., 2010; Fransson et al., 2007; Smyser et al., 2010; Peng et al., 2020). Other functional networks involved in heteromodal functions appear later. The DMN (Fox et al., 2007; Greicius et al., 2003; Greicius et al., 2009; Raichle et al., 2001; Raichle, 2015; Smith et al., 2009) is a higher-order functional network. Smyser et al (Smyser et al., 2010) found that SM and Vis functional networks mature earlier and demonstrate adult-like pattern for preterm neonate brain, with the DMN much immature and incomplete around birth. Cao et al (Cao et al., 2017a) also found rapid maturation of primary sensorimotor functional systems in preterm neonates from 31 to 41 postmenstrual weeks while the DMN remained immature during that period. These previous studies suggest that significant functional maturation in primary sensorimotor networks occur earlier in preterm and perinatal developmental period (Cao et al., 2017a; Doria et al., 2010; Smyser et al., 2010) compared to 0-24-month infancy focused in the present study. Functional network emergence in the DMN was found in the developmental infant cohort in Figure 3, marking significant maturation of the DMN in infancy and distinguished network pattern from earlier developmental period. The delineated DMN emergence in this study is also consistent to the literature (Gao et al., 2009). Figure 3-figure supplement 1 further demonstrated significant increase of FC only in the DMN, but not in primary sensorimotor system that already emerged in earlier developmental period.

Glucose and oxygen are two primary molecules for energy metabolism in the brain (Raichle et al., 2001; Vaishnavi et al., 2010). Glucose consumed by infant brain represents 30% total amount of glucose (Raichle, 2010; Settergren et al., 1976), more than 15-20% typically seen in adult brain (Bouma and Muizelaar, 1990; Satterthwaite et al., 2014). The cerebral metabolic rate for glucose (CMRGlu) and oxygen (CMRO2) are direct measures of the rate of energy consumption, which parallel the proliferation of synapses in brain during infancy (Raichle, 2010). RCBF delivering glucose and oxygen for energy metabolism in the brain is closely related to CMRGlu and CMRO2 and can serve as a surrogate of these two measurements (Fox and Raichle, 1986; Gur et al., 2009; Paulson et al., 2010; Vaishnavi et al., 2010). In the PET study (Chugani and Phelps, 1986) using CMRGlu measurements, it was found that the local CMRGlu in the sensorimotor cortex almost reaches the highest level in early infancy and then plateaus during rest of infancy, consistent with relatively small changes of rCBF in later infancy in primary sensorimotor ROIs found in this study (Figure 2). On the other hand, the global CBF measured with PC MRI (Liu et al., 2019) increases dramatically during infancy with global CBF at 18 months almost 5 times of the CBF around birth. Taken together, the literature suggests significant but nonuniformly distributed CBF increases across the brain regions during infancy, consistent to the measured heterogeneous rCBF increase pattern (Figures 1 and 2) in the present study.

Furthermore, the differentiated cerebral metabolic pattern reflected by measured rCBF distribution (Figure 1) at a specific age and differential increase rates of the rCBF (Figure 2) from 0 to 24 months are strikingly consistent with spatiotemporally differentiated functional (Cao et al., 2017a; Doria et al., 2010; Fransson et al., 2007; Gao et al., 2009; Smyser et al., 2010) and structural (Ouyang et al., 2019a) maturational processes. In developmental brains, cellular processes supporting differential functional emergence require extra oxygen and glucose delivery through cerebral blood flow to meet the metabolic demand. In Hebb’s principle, “neurons that fire together wire together”. Through the synaptogenesis in neuronal maturation, the neurons within a certain functional network system tend to have more synchronized activity in a more mature stage than in an immature stage. As shown in the diagram in Figure 6, cellular activities in developmental brains, such as synaptogenesis critical for brain circuit formation, need extra energy more than that in the stable and matured stage. Neurons do not have internal reserves of energy in the form of sugar or oxygen. The demand of extra energy requires an increase in rCBF to deliver more oxygen and glucose for formation of brain networks. In the context of infant brain development, there is a cascade of events of CBF increase, CMRO2 and CMRGlu increase, synaptogenesis and synaptic efficacy increase, blood oxygenation level dependent (BOLD) signal synchronization increase, and functional connectivity increase, shown in the bottom of Figure 6. Such spatial correlation of rCBF to the FC in the functional network ROIs was found in adults (Liang et al., 2013). Higher rCBF has also been found in the DMN in children 6 to 20 years of age (Liu et al., 2018). Consistent with the diagram shown in Figure 6, Figure 4 revealed significant correlation between FC and rCBF in the DMN network, and Figure 5 identified this significant correlation between FC and rCBF specifically in the DMN network, but not in the primary Vis or SM networks. Collectively, these results (Figures 4 and 5) were well aligned with the hypothesis that faster rCBF increase in the DMN underlies emergence of the DMN reflected by significant FC increases. The revealed physiology-function relationship may shed light on physiological underpinnings of brain functional network emergence.

**Figure 6.**
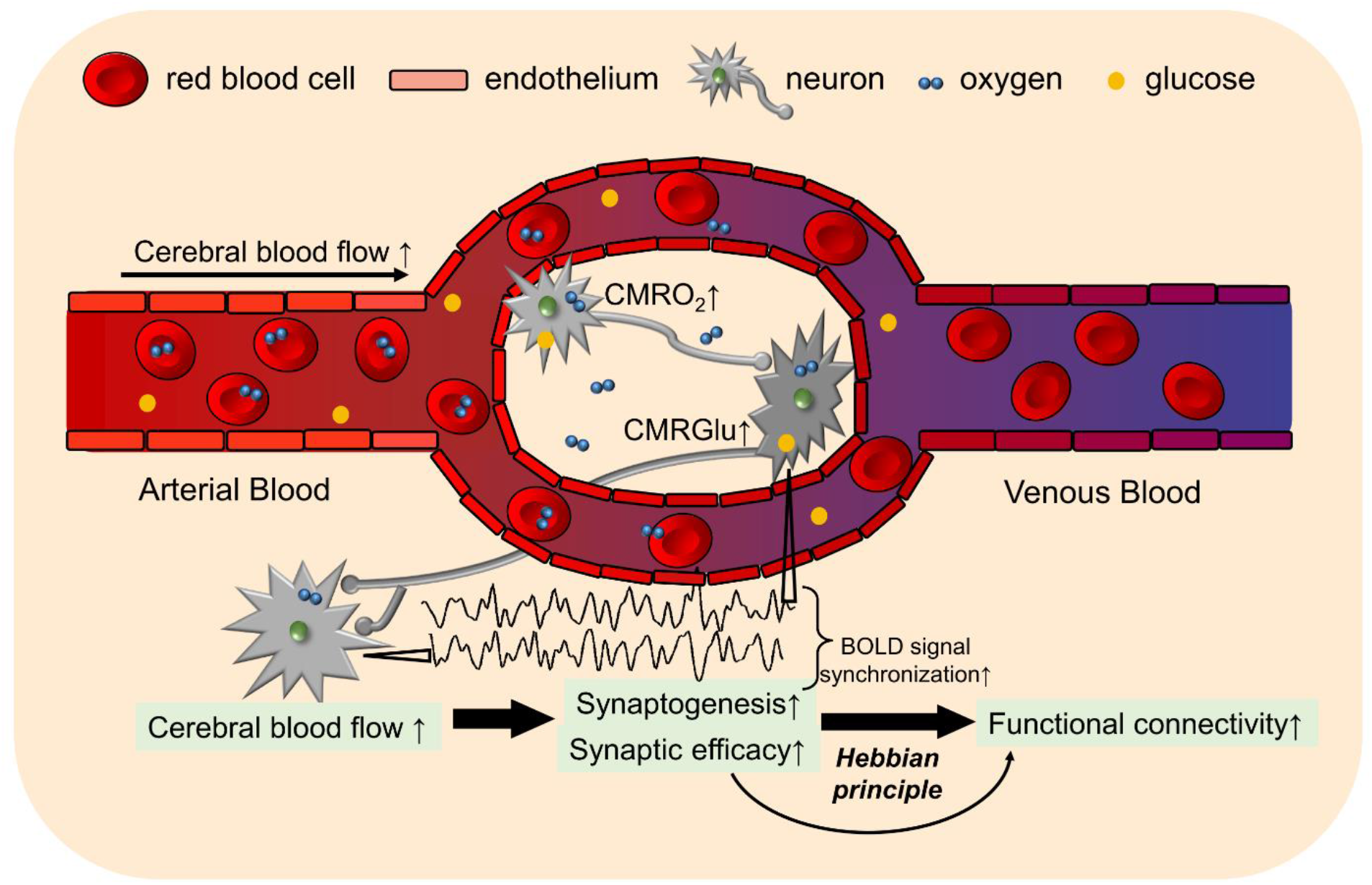
A diagram illustrating hypothesized neuronal mechanism supporting coupling of regional cerebral blood flow (rCBF) and functional connectivity. Specifically, higher cerebral blood flow delivers larger amount of oxygen and glucose to neurons, leading to cascade of events of regional cerebral metabolic rate of oxygen (CMRO2) ↑ and glucose (CMRGlu) ↑, synaptogenesis ↑ and synaptic efficacy ↑, blood oxygenation level dependent (BOLD) signal synchronization ↑, and functional connectivity ↑, during infant brain development.

It is noteworthy to highlight a few technical details below. First, present study benefits from multimodal MRI allowing measuring functional network emergence and rCBF of the same cohort of infants. Simultaneous rCBF and FC measurements enabled us to probe relationship of brain physiology and function during infant development. Second, a nonparametric permutation analysis without a prior hypothesis at the voxel level across the whole brain was conducted to confirm the coupling of rCBF and FC is specific in the DMN regions (Figure 5), not in the primary sensorimotor (Vis or SM) regions (Figure 5-figure supplement 2), demonstrating the robustness of the results on physiological underpinning of functional network emergence in the DMN. Third, the pCASL perfusion MRI was calibrated by PC MRI so that the errors caused by varying labeling efficiency among infant subjects can be ameliorated. This calibration process therefore enhanced accuracy of rCBF measurements for their potential use as a standard reference. Fourth, the ROIs for rCBF measurements were obtained by data-driven ICA analysis of the same sub-cohort of infant subjects aged 12 to 24 months instead of transferred ROIs from certain brain parcellation atlases. Since most of parcellation atlases were built based on adult brain data and all these atlases were established based on other subject groups, ROIs delineated from the same cohort improve accuracy of coupling analysis. There are several limitations in the present study that can be improved in future investigations. All data were acquired from a cross-sectional cohort. To minimize the inter-subject variability, future study with a longitudinal cohort of infants is warranted. With relatively small size of infant brains, spatial resolution of the pCASL and rs-fMRI can be further improved too to improve imaging measurement accuracy. To further improve the statistical power, larger infant sample size will be beneficial in the future studies. Finally, although the rCBF from pCASL perfusion MRI is highly correlated with the CMRO2 and CMRGlu measured from PET, rCBF is not direct measurement of rate of energy consumption. Physiology-function relationship studies in infants could benefit from development of novel noninvasive MR imaging methods as an alternative to PET to measure CMRO2 or CMRGlu without involvement of radioactive tracer.

## CONCLUSION

Novel findings in this study inform a physiological mechanism of DMN emergence during infancy with rCBF and FC measurements from multimodal MRI in developmental infant brains. The age-specific whole-brain rCBF maps and spatiotemporal rCBF maturational charts in all brain regions serve as a standardized reference of infant brain physiology for precision medicine. The rCBF-FC coupling results revealing fundamental physiology-function relationship have important implications in altered network maturation in developmental brain disorders.

## MATERIALS AND METHODS

### Infant subjects

Forty-eight infants (30 males) aged 0-24 months (14.6 ± 6.32 months) were recruited at Beijing Children’s Hospital. These infants were referred to MR imaging due to seizures with fever (*n* = 21), convulsion (*n* = 17), diarrhea (*n* = 9), or sexual precocity (*n* = 1). All infants had normal neurological examinations documented in medical record. The exclusion criteria include known nervous system disease, or history of neurodevelopmental or systemic illness. Every infant’s parents provided signed consent and the protocol was approved by Beijing Children’s Hospital Research Ethics Committee (Approval number 2016-36).

### Data acquisition

All infant MR scans including pCASL, PC MRI, rs-fMRI and structural MRI were acquired with the same 3T Philips Achieva system under sedation. Earplug and headphones were used to minimize noise exposure. Pseudo-continuous arterial spin labeling (pCASL) perfusion MRI images were acquired using a multi-slice echo planar imaging with following parameters: TR = 4100 ms, TE = 15 ms, 20 slices with 5 mm slice thickness and no gap between slices, field of view (FOV) = 230 × 230 mm^2^, matrix size = 84 × 84, voxel size = 2.74 × 2.74 × 5 mm^3^. As shown on the left panel of Figure 1a, the labeling slab was placed at the junction of spinal cord and medulla (65 mm below central slab of imaging volume) and parallel to the anterior commissure-posterior commissure (AC-PC) line. The labeling duration was 1650 ms and the post labeling delay (PLD) was 1600 ms. 30 pairs of control and label volume were acquired for each infant. The acquisition time of the pCASL perfusion MRI images was 4.2 min. An auxiliary scan with identical readout module to pCASL but without labeling was acquired for estimating the value of equilibrium magnetization of brain tissue. Phase contrast (PC) MRI was acquired to calibrate regional cerebral blood flow (rCBF) by scaling the overall CBF of entire brain. The carotid and vertebral arteries were localized based on a scan of time-of-flight (TOF) angiography acquired with following parameters: TR = 20 ms, TE = 3.45 ms, flip angle = 30°, 30 slices, FOV= 100 × 100 mm^2^, matrix size = 100 × 100, voxel size = 1 × 1 × 1 mm^3^. The acquisition time of the TOF angiography was 28 s. Based on the angiography, the slices for the PC MRI of internal carotid arteries were placed at the level of the foramen magnum and the slices for the PC MRI of vertebral arteries were placed between the two turns in V3 segments, as illustrated on the right panel of Figure 1a. For left and right internal carotid and vertebral arteries, PC MRI images were acquired with following parameters: TR = 20 ms, TE = 10.6 ms, flip angle = 15°, single slice, FOV = 120 × 120 mm^2^, matrix size = 200 × 200, voxel size = 0.6 × 0.6 × 3 mm, maximum velocity encoding= 40 cm/s, non-gated, 4 repetitions. The acquisition time of PC MRI for each artery was 24 s. Resting state fMRI (rs-fMRI) images were acquired using echo planar imaging with following parameters: TR = 2000 ms, TE = 24 ms, flip angle = 60°, 37 slices, FOV = 220 × 220 mm^2^, matrix size = 64 × 64, voxel size = 3.44 × 3.44 × 4 mm^3^. 200 dynamics were acquired for each infant. The acquisition time of the rs-fMRI images was 7 min. T1 weighted images of all infants were also acquired for anatomical information and brain segmentation using MPRAGE (Magnetization Prepared - RApid Gradient Echo) sequence with following parameters: TR = 8.28 ms, TE = 3.82 ms, flip angle = 12°, 150 slices, FOV = 200 × 200 mm^2^, matrix size = 200 × 200, voxel size = 1 × 1 × 1 mm^3^. The acquisition time of the T1-weighted image was 3.7 min. Visual inspection was carefully conducted for all MRI data by experienced pediatric radiologists (D.H. and Y.P.) with decades of experience in clinical radiology. No significant motion artifacts were spotted with the sedated MRI scans. Therefore, no dataset from the forty-eight infants was excluded from following data analysis due to severe motion artifacts.

### Measurement of rCBF with pCASL perfusion MRI and calibrated by PC MRI

After head motion correction of the pCASL perfusion MRI, we estimated rCBF using the protocol similar to that in our previous publication (Ouyang et al., 2017). Briefly, rCBF was measured using a model described in ASL white paper (Alsop et al., 2015):

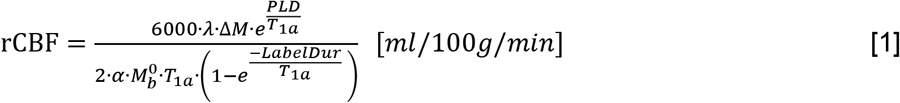

where ΔM is the dynamic-averaged signal intensity difference between in the control and label images; λ, the blood-brain partition coefficient, is 0.9 ml/g (Herscovitch and Raichle, 1985); PLD, the post labeling delay time, is the cumulation of 1650 ms and the delayed time between slices; LabelDur, the labeling duration, is 1600 ms; α, the labeling efficiency, is 0.86 predicted by the fitting between labeling efficiency and blood velocity in the previous study (Aslan et al., 2010); T_1a_, T_1_ of arterial blood, is 1800 ms (Liu et al., 2016; Varela et al., 2011). The value of equilibrium magnetization of brain tissue 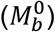 was obtained from an auxiliary scan with identical readout module of pCASL except labeling. The labeling efficiency α can vary considerably across participants, especially in infants. Thus, we used PC MRI to estimate and calibrate rCBF measures, as described previously (Aslan et al., 2010; Ouyang et al., 2017). To calibrate rCBF, global CBF from PC MRI was calculated as follows:

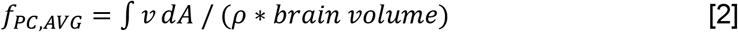

where v is the blood flow velocity in the ICAs and VAs; A is the cross-sectional area of the blood vessel with the unit mm^2^; and the brain tissue density ρ is assumed as 1.06 g/mL (Dittmer and Dawson, 1961; Herscovitch and Raichle, 1985). Brain volume was measured from the T1-weighted image as parenchyma volume (gray matter + white matter volume). RCBF was calibrated by applying the scalar factor making averaged rCBF equal to global CBF from PC MRI. To demonstrate the reproducibility of the adopted pCASL protocol, the intraclass correlation coefficient (ICC) was calculated based on entire brain rCBF maps measured from first half and second half of control/label series of pCASL scan of a randomly selected infant subject aged 17.6 months.

### Rs-fMRI preprocessing

Same preprocessing procedures elaborated in our previous publication (Cao et al., 2017) was used. Briefly, the normalized rs-fMRI images underwent spatially smoothing with a Gaussian kernel (full width at half-maximum of 4mm), linear trend removal, and temporal band-pass filtering (0.01–0.10 Hz). Several nuisance variables, including 6 rigid-body head motion parameters and the averaged signal from white matter and cerebrospinal fluid (CSF) tissue, were removed through multiple linear regression analysis to reduce effects of non-neuronal signals. The CSF and white matter were segmented with the T1-weighted image using SPM. Preprocessed rs-fMRI signals were used to estimate functional connectivity (FC), defined as Pearson’s correlation between the time courses of preprocessed rs-fMRI blood oxygenation level dependent (BOLD) signal in two regions or two voxels (Figure 1-figure supplement 2a).

### Multi-modal image registration to a customized template space from all subjects

For integrating perfusion MRI and rs-fMRI data of all infant subjects, a customized structural template was generated. T1-weighted image of a 12-month-old brain characterized by median brain size at this age and straight medial longitudinal fissure was used as a single-subject template. T1-weighted images of all subjects were registered to the single-subject template by using nonlinear registration in Statistical Parametric Mapping (SPM 8, http://www.fil.ion.ucl.ac.uk/spm). The averaged T1-weighted image in the template space was defined as the structural template. In individual space, after slice timing and head motion correction, intra-subject registration of rs-fMRI to T1-weighted image was conducted by transforming the averaged images across dynamics in each session to the T1-weighted image of the same subject through linear registration with SPM. This intra-subject transformation was applied to each volume of rs-fMRI. As described in the main text, rCBF map was estimated in the individual space using Eq [1]. Intra-subject registration of rCBF map to T1-weighted image was also conducted through linear registration with SPM. RCBF maps and rs-fMRI images aligned to T1-weighted images in the individual space of each subject were then normalized to the customized infant template space through the same nonlinear registration from individual T1-weighted image to the structural template. After nonlinear inter-subject normalization, rCBF map was smoothed spatially in the customized template space.

### Identification of functional network regions-of-interests (ROIs) with rs-fMRI

Brain regions of three functional networks, including default-mode network (DMN), visual (Vis) network, and sensorimotor (SM) network, were used as functional ROIs for quantifying rCBF. These functional networks were identified with rs-fMRI of 35 infants aged 12 to 24 months, as DMN can be better delineated in later infancy than earlier infancy. Independent Component Analysis (ICA) (Beckmann et al., 2005) in the FSL (https://fsl.fmrib.ox.ac.uk/fsl/fslwiki/MELODIC) was used to identify all these network regions. Individual DMN regions, including posterior cingulate cortex (PCC), medial prefrontal cortex (MPFC), inferior parietal lobule (IPL) and lateral temporal cortex (LTC), were extracted from the independent components (ICs) based on the spatially distributed regions consistently identified across studies (Greicius et al., 2004; Smith et al., 2009). Vis and SM network regions were also identified following the literature (Smith et al., 2009). The functional network ROIs were obtained by thresholding the ICs at z value of 1.96 which corresponds to p value of 0.05. The identified ROIs for DMN, Vis and SM networks are shown in Figure 1-figure supplement 2b. ROIs of DMN, Vis or SM include all regions in each network, respectively.

### Characterization of age-dependent changes of rCBF and FC

After multi-modal images from all subjects were registered in the same template space, age-dependent rCBF and FC changes were characterized using linear regression. The linear cross-sectional developmental trajectory of voxel-wise rCBF increase was obtained by fitting rCBF measurements in each voxel with ages across subjects. The fitted rCBF at different ages was estimated and projected onto the template cortical surface with Amira (FEI, Hillsboro, OR) to show the spatiotemporal changes of rCBF across cortical regions during infancy. The rCBF-age correlation coefficients r values of all voxels were estimated and mapped to the cortical surface, resulting in the correlation coefficient map. The functional ROIs identified above were used to quantify regional rCBF change. The rCBF in the SM, Vis, and DMN ROIs were averaged across voxels within each ROI, and fitted with a linear model: rCBF(t) = α + β t + ε, where α and β were intercepts and slopes for rCBF measured at certain ROI, t was the infant age in months and ε was the error term. To test if regional rCBF increased significantly with age, the null hypothesis was that rCBF slope in each ROI was equal to zero. To compare regional rCBF change rates between the tested ROI and SM ROI, the null hypothesis was that the rCBF slope of the tested ROI and rCBF slope of SM ROI were equal. Rejection of the null hypothesis indicated a significant rCBF slope difference between two ROIs.

Nonlinear models, including exponential and biphasic models, were compared with linear model using F-test for fitting regional rCBF in the DMN, SM and Vis ROIs. No significant difference was found between linear and exponential (all F(1,45) < 3.45, p > 0.05, uncorrected) fitting or between linear and biphasic (all F(2,44) < 2.64, p > 0.05, uncorrected) fitting in the regional rCBF changes in the DMN, SM and Vis ROIs.

In the customized template space, time-dependent changes of FC to PCC and time-dependent changes of FC within the functional network ROIs were calculated. FC of any given voxel outside the PCC to PCC was defined as correlation between signal time course of this given voxel and the averaged signal time course of all voxels within the PCC. Emergence of the DMN was delineated by linear fitting of the age-related increase of FC to PCC in each voxel across subjects. FC within each network was defined as mean of functional connectivity strength (Cao et al., 2017b) of each voxel within this network region. The FC within the DMN, SM or Vis was also fitted with a linear model: FC(t) = α + β t + ε, where α and β were intercepts and slopes for FC measured within a certain network, t was the infant age in months and ε was the error term. To test if the FC within a network increased significantly with age, the null hypothesis was that rCBF slope in each ROI was equal to zero.

### Test of heterogeneity of rCBF across functional network ROIs

To examine heterogeneity of infant rCBF in different brain regions, rCBF measurements of all infants were averaged across voxels in the Vis, SM and DMN (including DMN subregions DMN_PCC, DMN_MPFC, DMN_IPL and DMN_LTC), respectively. To test significant difference of the rCBF values among different functional network ROIs, a one-way analysis of variance (ANOVA) with repeated measures was conducted. Paired t-tests were also conducted to test the difference of rCBF measurements between regions. False discovery rate of each test was corrected to control the type I error.

### Coupling between rCBF and FC during the infant brain development

Coupling between rCBF and FC in the DMN was conducted with voxel-wise approach. The FC of a voxel in the DMN was the average of correlations of rs-fMRI BOLD signal between this voxel and all other DMN voxels. All infants were divided into two groups based on their ages, 0-12 months and 12-24 months. 4000 voxels were randomly chosen from the DMN voxels of all subjects in each age group for regression analysis. Since the variance of both FC and rCBF cannot be ignored in this study, Deming regression (Deming, 1943) was used to fit the trendline of coupling between FC and rCBF. We further tested whether significant FC-rCBF coupling was specifically localized in the DMN, but not in primary sensorimotor (Vis or SM) regions. FC within a specific network was calculated by averaged FC of all voxels in this network. As demonstrated in Figure 5-figure supplement 1, the correlation between FC within a network and the rCBF at each voxel resulted in a whole-brain r map (e.g. Figure 5b). A nonparametric permutation test was then applied to evaluate the significance of rCBF-FC correlation in the ROIs of a specific brain network (e.g. DMN, SM or Vis). The null hypothesis is that the voxels with significant correlation (*r* > 0.28) between rCBF and FC are distributed evenly in the brain. To test the null hypothesis, we resampled the correlation coefficient r of all brain voxels randomly for 10,000 times to build 10,000 whole-brain correlation coefficient distribution maps. The FC-rCBF correlation was considered significant in certain brain network ROIs if the number of observed significant voxels in the network ROIs is higher than the number of significant voxels corresponding to 95th percentile in the permutation tests.

## ACKNOWLEDGEMENTS

This work was supported by grants from National Institute of Health: R01MH092535, R01MH125333, R01EB031284, R21MH123930 and P50HD105354.

## AUTHOR CONTRIBUTIONS

H.H. designed the research; Q.Y., M.O., H.K., D.H., Y.P. and H.H. performed research; Q.Y. and H.H. contributed new reagents/analytic tools; Q.Y., M.O., J.D., B.H., F.F. and H.H. wrote the paper.

## Figures and figure supplements

**Figure 1-figure supplement 1.**
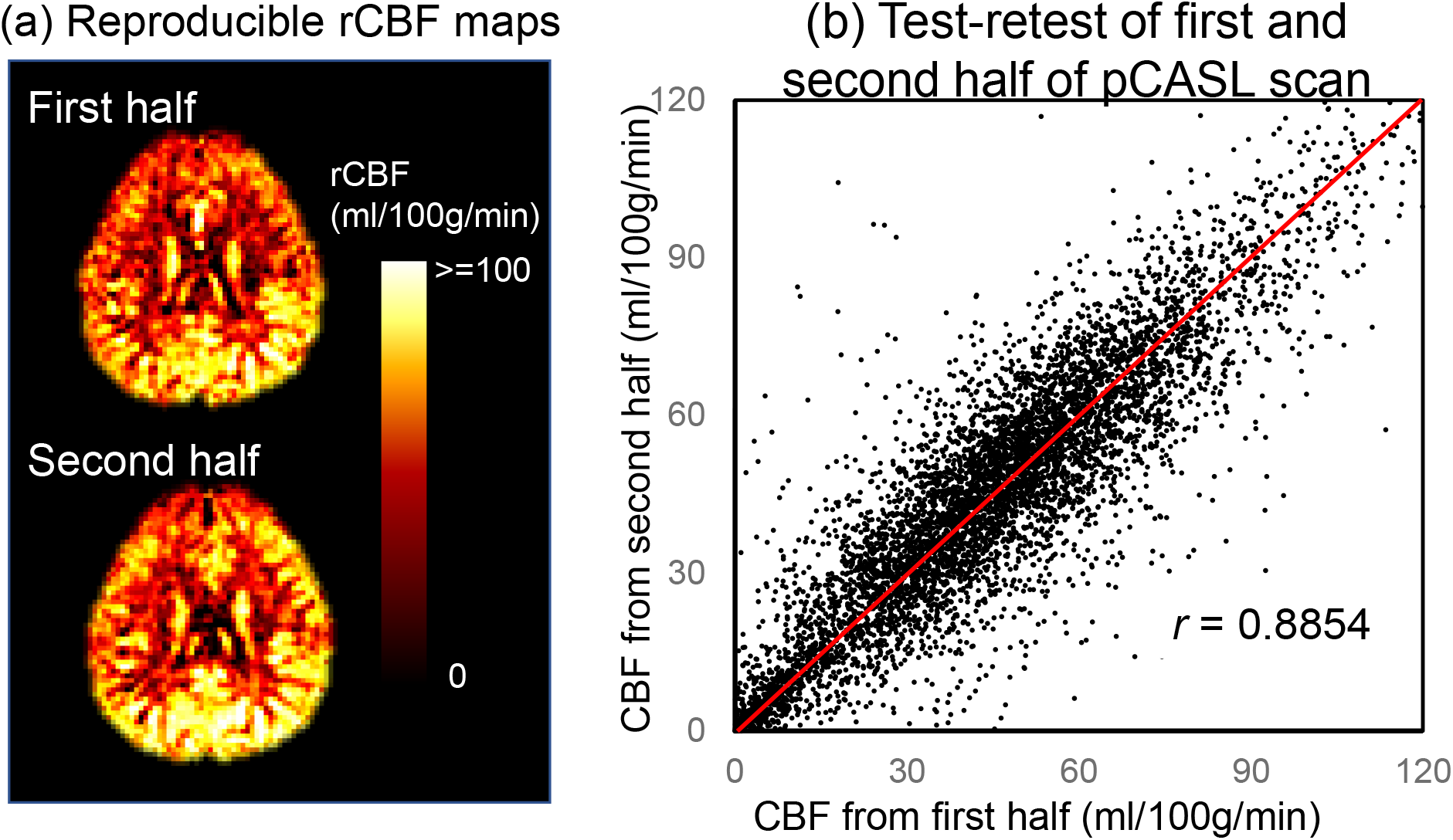
Highly reproducible pCASL protocol adopted in the present study for measuring rCBF. Panel (a) shows reproducible rCBF maps measured from first and second half of the control/label volumes, and panel (b) shows the intraclass correlation coefficient (ICC) 0.8854 with the 95% confident intervals of [0.88, 0.8906] calculated from entire brain rCBF maps measured from first and second half control/label volumes. Both panels (a) and (b) are generated based on pCASL scans of a randomly selected infant subject aged 17.6 months.

**Figure 1-figure supplement 2.**
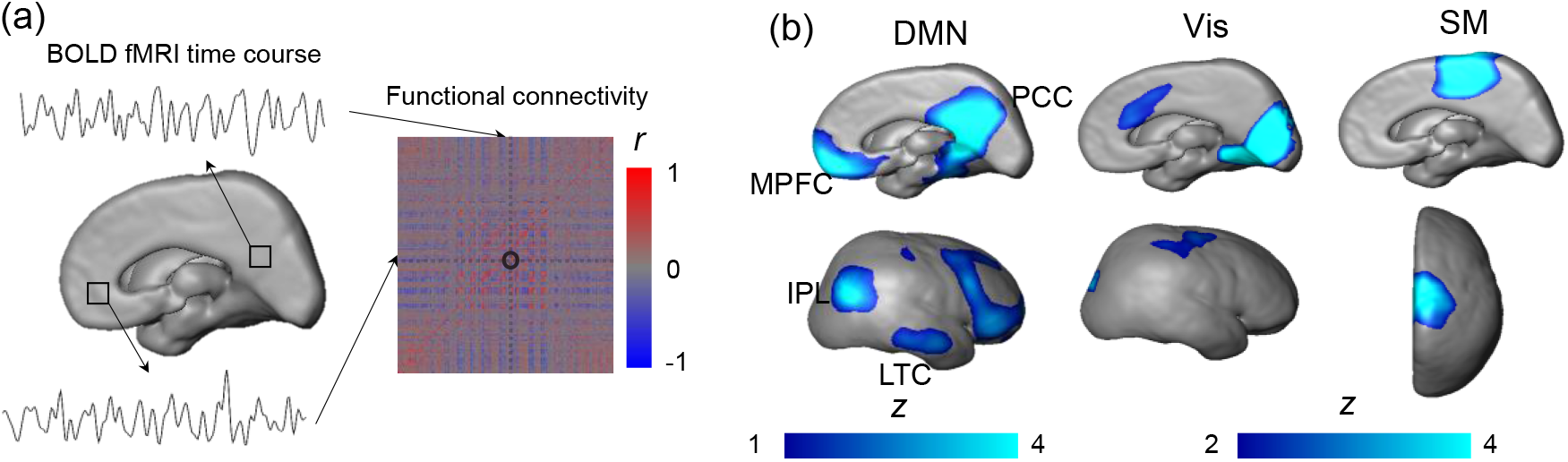
Identification of functional network regions-of-interests with resting-state fMRI. (a) Functional connectivity obtained from correlation between blood oxygenation level dependent (BOLD) fMRI time courses of two voxels; (b) Brain regions of the visual (Vis), sensorimotor (SM) and default-mode network (DMN) identified by z statistic maps from group independent component analysis. DMN subregions posterior cingulate cortex (PCC), medial prefrontal cortex (MPFC), inferior posterior lobule (IPL) and lateral temporal cortex (LTC) can be clearly appreciated.

**Figure 1-figure supplement 3.**
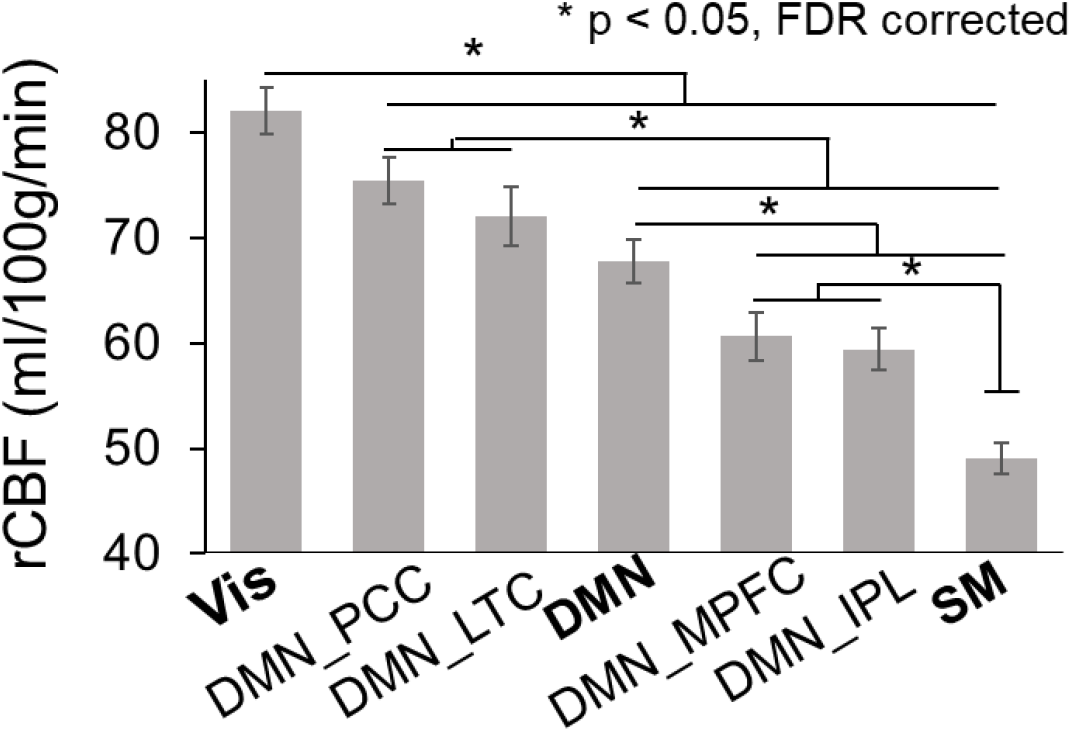
Heterogeneity of rCBF measurements across functional network regions. Significant differences (*p* < 0.05 with false discovery rate (FDR) correction) of rCBF measurements between regions were indicated by asterisks.

**Figure 3-figure supplement 1.**
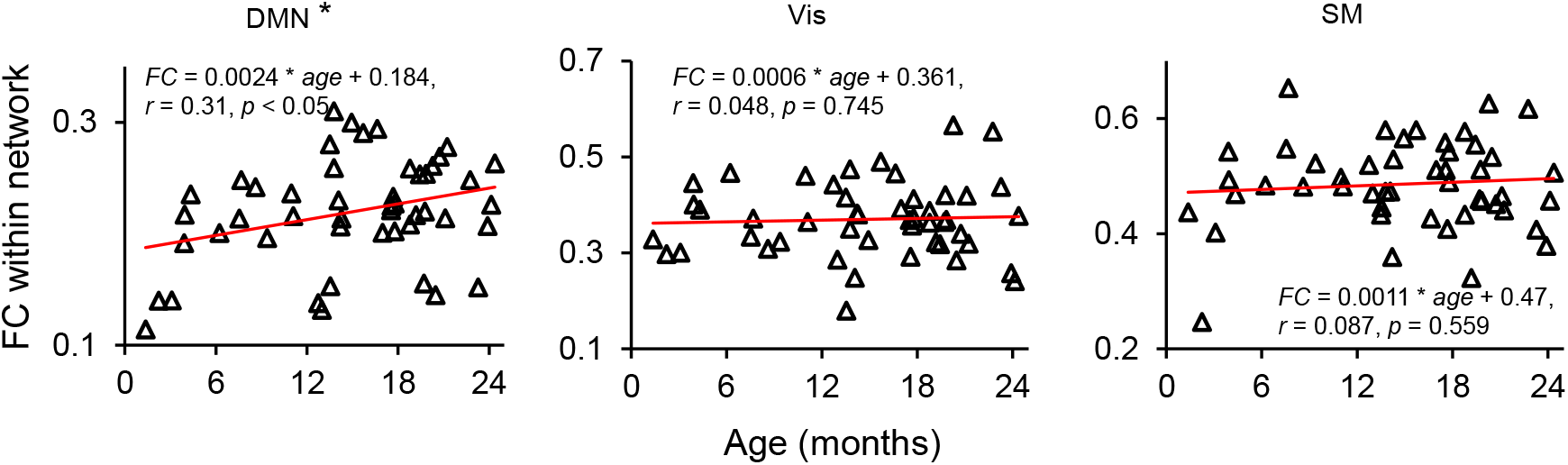
Age-dependent changes of functional connectivity (FC) within the default-mode network (DMN), visual (Vis) and sensorimotor (SM) network regions during infancy. Significant age-dependent increase of within-network FC was found only in the DMN (*r* = 0.31, *p* < 0.05, indicated with an asterisk) regions, but not in the Vis (*r* = 0.048, *p* = 0.745) and SM (*r* = 0.087, *p* = 0.559) regions.

**Figure 5-figure supplement 1.**
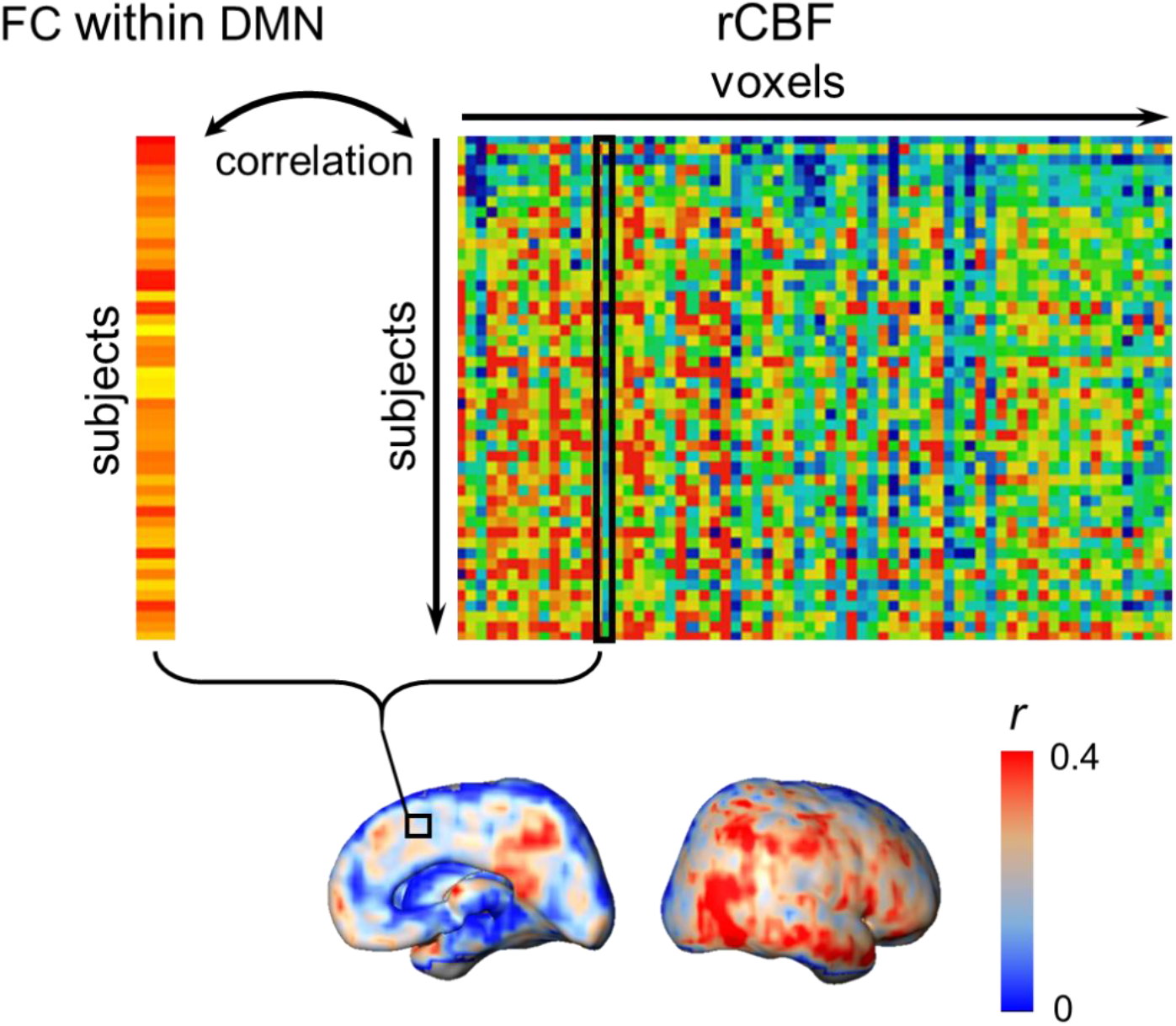
Procedures of generating map of correlation between the default-mode network (DMN) functional connectivity (FC) and reginal cerebral blood flow (rCBF) across the entire brain voxels. At any brain voxel indicated by a small black box in the correlation map below, correlation between the DMN FC across subjects and the rCBF at this voxel across subjects was conducted and the correlation coefficient was calculated for this voxel. The correlation map between the DMN FC and rCBF across the brain can be generated by projecting the correlation coefficient *r* of each voxel onto the cortical surface.

**Figure 5-figure supplement 2.**
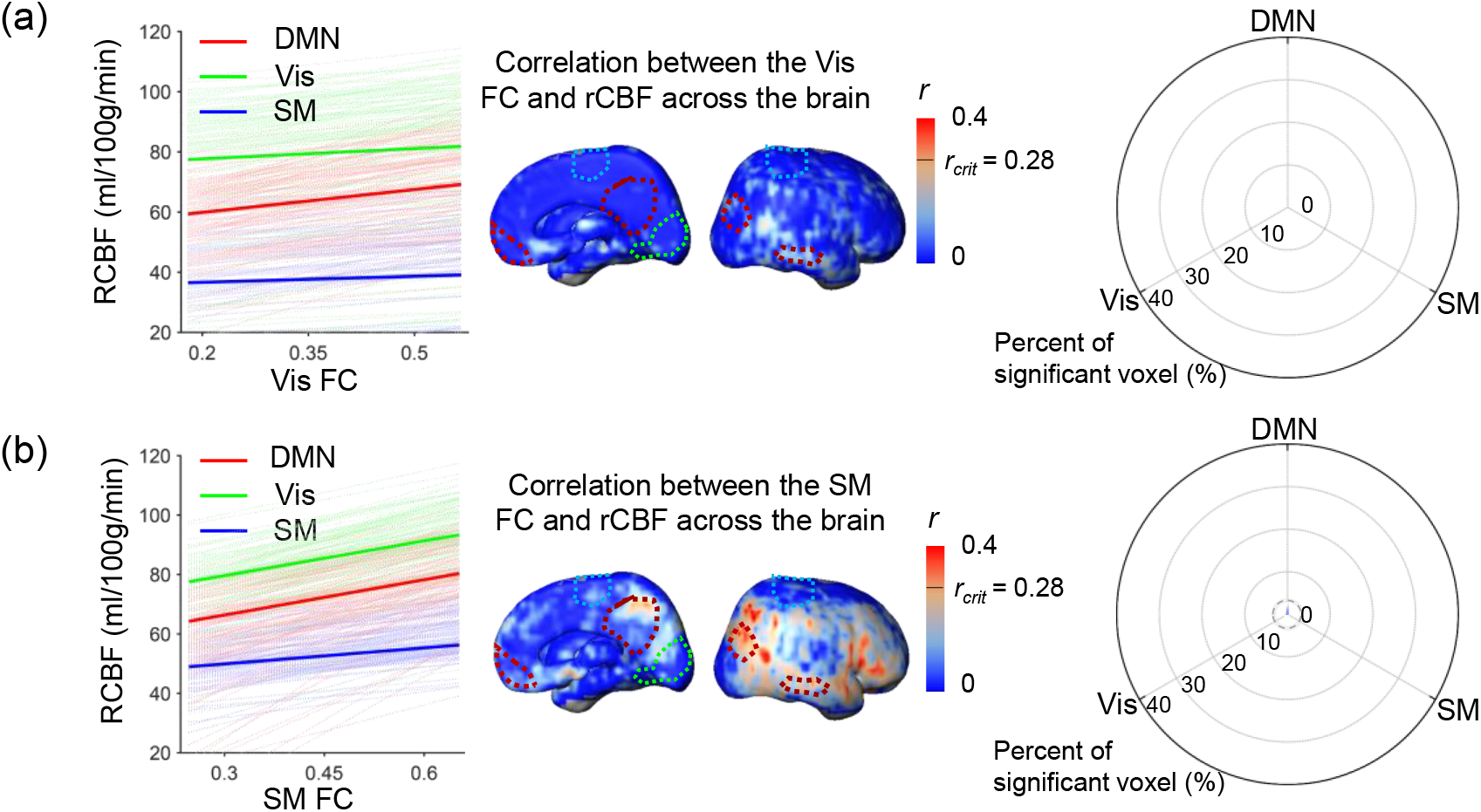
Coupling between intra-visual network functional connectivity (Vis FC) and regional cerebral blood flow (rCBF) and coupling between intra-sensorimotor network (SM FC) and rCBF. Left panels show the correlation of Vis FC (a) or SM FC (b) and rCBF at randomly selected voxels in the default-mode (DMN) (light red lines), Vis (light green lines) and SM (light blue lines) network regions. Correlations of Vis FC (a) or SM FC (b) and averaged rCBF in the DMN, Vis and SM network regions are shown as thickened red, green and blue lines, respectively. Middle panels demonstrate the distribution of voxel-wise correlation coefficient (r) obtained from correlation between Vis FC (a) or SM FC (b) and rCBF at each voxel. The short black line in the color bar indicates critical correlation coefficient r value rcrit corresponding to p=0.05. Low r values are apparent in all network (DMN, Vis and SM) regions with dashed red, green and blue contours (from Supplementary Fig. 2b) delineating DMN, Vis and SM network, respectively. Right panels show radar plot with radius representing percent of voxels with significant correlation between Vis FC or SM FC and rCBF in the DMN, Vis or SM network regions. The radii in radar plots demonstrate that the percent of voxels with significant correlations between intra-Vis (a) or intra-SM (b) FC and rCBF in DMN, Vis or SM network regions is 0 or close to 0, indicating that coupling between Vis FC or SM FC and rCBF in any network region is not significant. The dashed line circle in the radar plot indicates critical percent of significant voxels with p=0.05 from 10,000 permutation tests.

**Video 1:** Video of the 4D spatiotemporal whole-brain dynamics of regional cerebral blood flow from 0 to 24 months.

